# Aging impairs primary task resumption and attentional control processes following interruptions

**DOI:** 10.1101/2021.12.20.473604

**Authors:** Marlene Rösner, Bianca Zickerick, Melinda Sabo, Daniel Schneider

## Abstract

Attentional selection of working memory content is impaired after an interruption. This effect was shown to increase with age. Here we investigate how electrophysiological mechanisms underlying attentional selection within working memory differ during primary task resumption between younger and older adults. Participants performed a working memory task, while being frequently interrupted with either a cognitively low- or high-demanding arithmetic task. Afterwards, a retrospective cue (retro-cue) indicated the working memory content required for later report. The detrimental effect of the interruption was evident in both age groups, but while younger adults were more strongly affected by a high-than by a low-demanding interruption, the performance deficit appeared independently of the cognitive requirements of the interruption task in older adults. A similar pattern was found regarding frontal-posterior connectivity in the theta frequency range, suggesting that aging decreases the ability to selectively maintain relevant information within working memory. The power of mid-frontal theta oscillations (4-7 Hz) featured a comparable effect of interruptions in both age groups. However, posterior alpha power (8-14 Hz) following the retro-cue was more diminished by a preceding interruption in older adults. These results suggest an age-related deficit in the attentional selection and maintenance of primary task information following an interruption that appeared independent from the cognitive requirements of the interrupting task.

## 1 Introduction

Our everyday life is characterized by a high number of interferences, such as interrupting tasks (González & Mark, 2004). Importantly, the performance deficit as a result of such task interruptions occurs to a greater extent for older adults (Clapp et al., 2011; Mishra et al., 2013; Solesio-Jofre et al., 2011). While studies point to multiple age-related cognitive deficits (Cabeza & Dennis, 2013), it is also emphasized that these deficits come into play primarily when a task requires the attentional suppression of irrelevant information (e.g. Gazzaley et al., 2008; Hasher & Zacks, 1999). It can thus be suggested that the age-related deficit in dealing with interruptions is largely due to less efficient attentional control processes required for switching between a primary and an interrupting task. The current study therefore uses oscillatory correlates of attentional control processes in the EEG to investigate the extent to which these very processes are impaired during the resumption of a primary task after an interruption in older compared to younger adults.

In general, interruptions can be defined as secondary tasks requiring a response and the intention to resume the primary task afterwards (Clapp et al., 2011; Monk et al., 2008; Trafton et al., 2011). Consequently, the interrupted person is forced to allocate attentional resources towards completing the interruption task and then reactivate information of the interrupted task (primary task) thereafter (Clapp & Gazzaley, 2012). In this regard, the short-term storage of information in working memory is of central importance, because it allows the flexible allocation of attention between mental representations of different tasks (Baddeley et al., 2011; de Vries et al., 2018; Rösner et al., 2020; Schneider et al., 2019). There are several reasons to assume that the ability to reallocate attention to working memory representations following an interruption will be even worse for older than for younger adults: Firstly, the ability to refresh recently presented information (i.e., to verbally repeat the word presented last) is impaired by aging (Johnson et al., 2002, 2004), which is related to reduced prefrontal cortex activity (Johnson et al., 2004; Mitchell et al., 2000). As the refreshing of task-relevant information reflects an important mechanism for primary task resumption (Clapp et al., 2010), it can be assumed that older adults may have problems refocusing their attention on working memory representations following interruptions. In line with the inhibitory deficit hypothesis (Hasher & Zacks, 1999), it has further been shown that aging is linked to an impaired ability to inhibit information no longer required for an ongoing task (e.g., Gazzaley, et al., 2008). Consequently, the refocusing of attention to a primary task could be compromised by a reduced ability to inhibit the no-longer relevant interruption information or to exclude it from ongoing storage in working memory (Clapp & Gazzaley, 2012).

Prior research on attentional control mechanisms on the level of working memory has proposed neural oscillations in the theta (~4-7 Hz) and alpha (~8-14 Hz) frequency band as reliable correlates of the switching of the focus of attention between different working memory representations. For example, posterior alpha power has been shown to decrease during the attentional selection of visual information stored in working memory (Sauseng et al., 2005; Schneider et al., 2017). Furthermore, an increase in oscillatory power in the theta frequency range (~4-7 Hz) has been observed when shifting the focus of attention between working memory representations of two different tasks (de Vries et al., 2018). Also, in the context of task interruptions, we were able to relate impaired attentional reallocation to the primary task to reduced posterior alpha power suppression and lower mid-frontal theta power (Zickerick et al., 2021). The present study therefore uses these oscillatory correlates of attentional control processes to investigate which cognitive mechanisms are impaired when dealing with interruptions in older age.

For this reason, the experimental paradigm required participants to perform different types of interruption tasks in-between the storage of two stimulus orientations in working memory. The resumption of the primary task was signaled by a retrospective cue (retro-cue) indicating which of the stored mental representation from the primary task had to be reported at the end of a trial. Before retro-cue presentation, participants were interrupted in half of the trials by either a cognitively high- or low-demanding arithmetic task. While previous studies on the influence of aging on the handling of interruptions have used working memory tasks with binary decisions (Clapp et al., 2011; Clapp & Gazzaley, 2012; Shum et al., 2013), the current task made use of a continuous report procedure. It thus provided the opportunity to examine how the accuracy of working memory content from the primary task is affected by interruptions, independent of the speed of the decision. In line with earlier research, we expect older adults to be more strongly affected by interruptions (Arnau et al., 2019; Clapp & Gazzaley, 2012). This should be reflected in a larger interruption-related deficit in terms of the accuracy in reporting the orientation of the cued stimulus from the primary task. The use of a retro-cue further allowed us to analyze age-related differences in oscillatory correlates of the attentional selection of primary task working memory representations following interruptions. We expected that an age-related deficit in attentional reallocation back to the primary task would be reflected in reduced frontal theta power and lower posterior alpha power suppression relative to conditions without interruption. As a further parameter, we calculated the phase-locking of theta oscillations (phase lag index or PLI; see Stam et al., 2007) between a fronto-central cluster of electrodes and other recording sites (Arnau et al., 2019; Capotosto et al., 2009; Greenberg et al., 2010; Hopfinger et al., 2000). Prior research has shown that the theta phase coupling between frontal and parieto-occipital sites is linked to the ability to actively maintain selected relevant information in working memory and is affected by age (Tóth et al., 2014; Werkle-Bergner et al., 2006). We thus expected a reduction in theta connectivity between frontal and posterior brain regions for older adults following the retro-cues. This age-related difference could occur especially after interruptions since these require the reactivation of primary task information in working memory.

In summary, this study will extend previous research on differences in primary task resumption following interruptions between younger and older adults by investigating neural oscillatory correlates of attentional control processes. We propose that respective age-related differences will be reflected in modulations of oscillatory parameters associated with attentional control processes to refocus on the primary task.

## 2 Methods

### 2.1 Participants

Twenty-five younger and 23 older adults took part in the experiment. Data from the younger participants came from a subsample of the experiment by Zickerick and Rösner, et al. (2021). Five older participants were excluded due to performance below chance level in the interruption task, leading to a final sample of 43 participants (25 younger adults: *M(age)* = 23.96, *SD* = 3.18, range = 19 - 29, 16 females; 18 older adults: *M(age)* = 67.06, *SD* = 4.7, range = 60 - 73, 10 females). All participants were right-handed according to a handedness questionnaire (adapted from Oldfield, 1971) and reported normal or corrected-to-normal vision. Color vision was confirmed with the Ishihara test for color blindness. Participants with neurological or psychiatric disorders (based on self-report) were excluded from participation.

Memory and executive functions in both age groups were tested with a short neuropsychological test battery (see table 1) prior to the preparation of the EEG recording. Older and younger adults did not statistically differ regarding their performance in the number repetition test included in the ‘Nürnberger Altersinventar’ (NAI) task that measures verbal working memory performance (Oswald & Fleischmann, 1999). However, selective attention and information processing speed was decreased in older compared to younger adults (trail making test or TMT; TMT-A and TMT-B). Furthermore, older adults were more susceptible to interference than younger adults, as assessed by color-word-test included in the NAI (Oswald & Fleischmann, 1999). Importantly, in the latter cases all older adults were still within the norm criteria for their age group. Thus, no subjects were excluded based on the neuropsychological testing.

**Table1.** Demographics of the sample and results of the neuropsychological assessment for younger and older adults

| Demographic and neuro-psychological variables | younger adults<br>(n = 25) | older adults<br>(n = 18) | *t(df)* |
| --- | --- | --- | --- |
| Sex (female/male) | 16/9 | 10/8 |  |
| Age(years) | 23.96 (3.2), 19-29 | 67.06 (4.8), 60-73 |  |
| Number repetition forward and backward | 12-72 (1.62), 10-16 | 11.94 (1.67), 9-15 | *t(41) = -1.53; p = 0.133* |
| TMT part A | 22.08 (5.37), 14-37 | 34.72 (8.33), 21-52 | *t(26) = 5.65; p < 0.001* |
| TMT part B | 48 (11.61), 28-73 | 82.94 (24.58), 51-126 | *t(22) = 5.60; p < 0.001* |
| Color-word-test | 7.28 (3.16), 2-14 | 18.82 (12.07), 7-59 | *t(17) = 3.86; p = 0.001* |

Participation was compensated with 10 € per hour or course credit. The study was approved by the local ethics committee of the Leibniz Research Centre for Working Environment and Human Factors. We furthermore considered the Declaration of Helsinki for medical research involving human subjects.

### 2.2 Apparatus, stimuli and, procedure

After EEG preparation, participants were seated in an electrically shielded, dimly lit chamber at 150 cm viewing distance to a 22-inch CRT monitor (100 Hz, 1024 x 768 pixels). Stimuli were generated with a ViSaGe MKII Stimulus Generator (Cambridge, Research Systems, Rochester, UK).

Prior to the beginning of the actual experiment, participants performed a training block on how to adjust the orientation of a visual stimulus by moving the computer mouse. Two randomly oriented bars (size: 1° by 0.1° visual angle) were presented to the left and right visual fields with 2.12° (visual angle) distance from a central fixation cross. Participants had to adjust the orientation of the grey bar (RGB: 140 140 140) to the orientation of the black bar (RGB: 0 0 0) by moving the computer mouse laterally. They clicked the left mouse button when they judged their orientation adjustment as complete. Participants were instructed to respond faster when the time required for adjusting the bar’s orientation exceeded 4000 ms. Additionally, the mean angular error had to be below 18° within a block of 50 trials. Participants had time to reach this criterion within an overall block of 150 trials. All participants were successful in this regard.

After the orientation adjustment training, participants performed two additional practice blocks. In the first one, they performed the primary task of the main experiment without interruption (28 trials). The second training block consisted of 36 trials in which participants performed the high-demanding interruption task (12 trials), the low-demanding interruption task (12 trials), and the prolonged fixation condition (12 trials) (see below for further information on the experimental paradigm).

In the actual experiment, a black fixation cross (RGB: 0 0 0, 1.7 ° x 1.7°) was presented in the center of the screen. In the initial memory array, two blue bars (RGB: 0, 176, 255, size: 0.1° x 1.0°, luminance: 52 cd/m^2^) were presented either on the left or right side of the screen for 200 ms (background RBG: 44, 44, 44, luminance: 25cd/m^2^; see figure 1). On the other side two grey sensory filler bars (luminance: 52 cd/m^2^, RGB: 140, 155, 126, size: 0.1° x 1.0°) were presented to prevent imbalanced sensory input with respect to luminance. Participants were instructed to memorize the orientation of the blue bars. After a delay interval of 800 ms with only the fixation cross present, a visual cue (size of the rectangle surrounding it: 3.44° x 2.58° visual angle) appeared in the center of the screen and indicated whether the primary task was going to be interrupted or not (see figure 1). A circle (size: 1.7° x 1,7°) indicated a prolonged presentation of the fixation cross for 2000 ms before the memory task was continued (prolonged fixation condition, ¼ of all trials). Two different types of interruptions were included in the experiment. A high-demanding interruption was indicated by a plus sign (size: 1.7° x 1.7°; high-demanding interruption condition, ¼ of all trials). During this interruption, participants were confronted with an arithmetic task presented on the center of the screen and had to judge whether this task was solved correctly by clicking either the left or right mouse button (index vs. middle finger). The math equation consisted of two one-digit numbers which always added up to a two-digit number to ensure comparable difficulty across trials. When the pre-cue was a minus sign (size: 1.7° x 1.7°), participants had to perform a low-demanding number comparison task (low-demanding interruption condition, ¼ of all trials). During the low-demanding interruption, two single-digit numbers, which always added up to a two-digit number, were presented as a vertical array in the center of the screen. Participants had to indicate by clicking either the left or right mouse button if the higher or lower number was presented at the bottom of the screen. Each interruption task had to be completed within 2000 ms. The assignment of response keys for the interruption tasks was counterbalanced across participants. The interruptions were followed by a 500 ms delay interval. Furthermore, a condition without any interruption or delay (early probe condition) was indicated by an “X” (size: 3.44° x 2.58°). This condition severed as a control condition to rule out the possibility that the performance differences between the interruption conditions and the prolonged fixation condition were due to a benefit resulting from a longer time interval available for working memory rehearsal in the prolonged fixation condition, instead of the detrimental effects related to the interruption itself. The early probe condition was not included in the EEG analysis.

**Figure 1.**
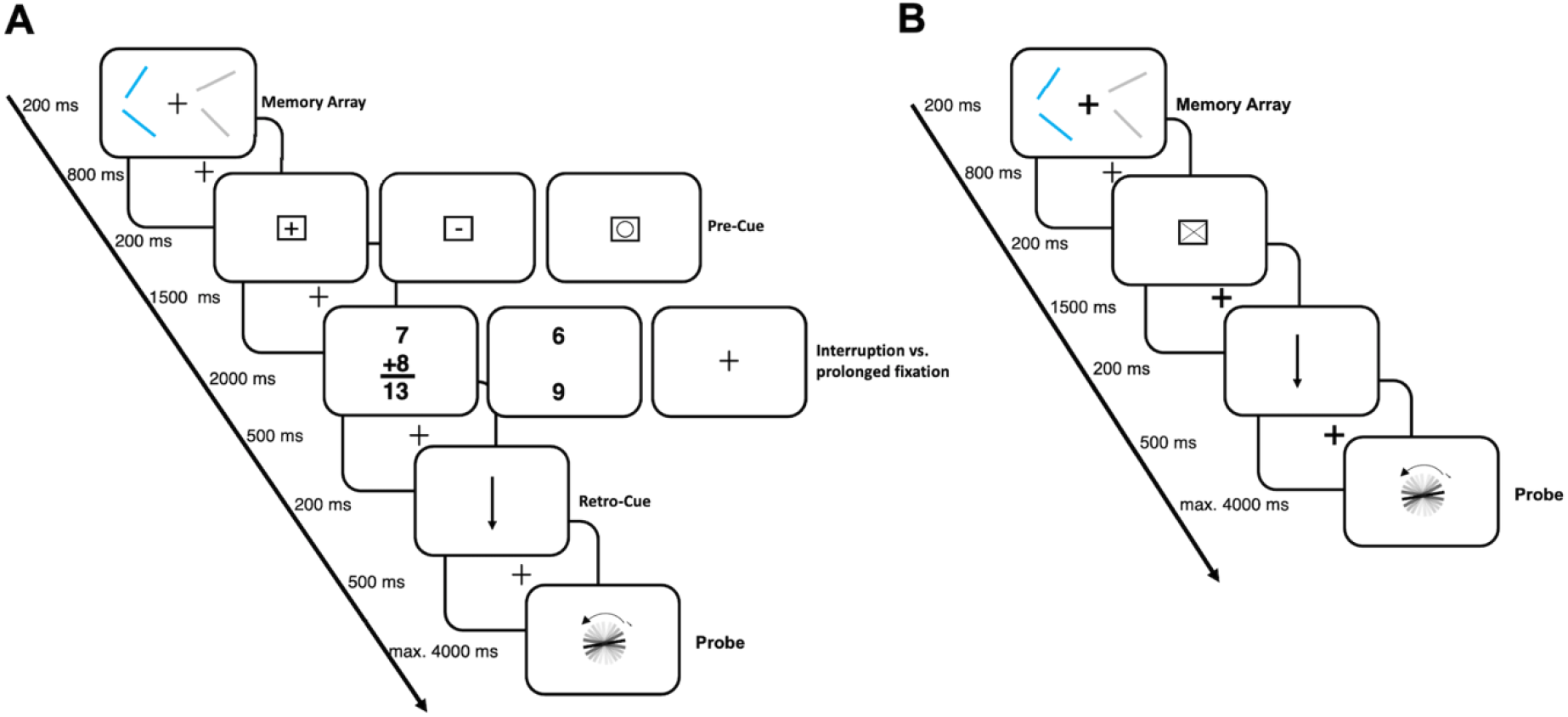
Experimental design. At the beginning of each trial, a memory array was presented consisting of two laterally presented blue bars with a random orientation and two task-irrelevant grey bars. To complete the working memory task, participants had to remember the blue bars’ orientation. At the end of each trial, a retro-cue indicated the bar whose orientation had to be reported. Panel A depicts the three conditions with either a high-demanding arithmetic task (decide whether the math equation is correct or false; high-demanding interruption condition), a low demanding number comparison task (decide whether the lower number is larger or smaller than the upper one; low-demanding interruption condition), or a prolonged fixation cross. The additional control condition, in which the orientations had to be stored over a short interval (retro-cue presentation at interruption task onset; early probe condition) is depicted in panel B.

Following the intervals with or without interruption, an arrow (retro-cue, size: 0.69° x 1.12° visual angle) was presented in the center of the screen indicating the position (top vs. bottom) of the stimulus whose orientation had to be reported at the end of the trial. A randomly oriented probe stimulus appeared in the center of the screen after another delay interval of 500 ms. Participants had to adjust the probe bar to the target orientation within 4000 ms and click the left button on the computer mouse when they judged their adjustment as complete. The probe stimulus remained visible for 200 ms following this response. The subsequent inter-trial interval varied randomly in steps of 1 ms between 500 and 1000 ms.

Participants performed 480 trials in total (120 trials per condition). Trials were separated into six blocks by breaks of about two minutes to prevent fatigue during the experiment. A trial of the early probe condition took about 5.5 s while each trial in the other conditions took about 7.9 s. Including the neuropsychological testing and the EEG preparation, the whole experiment lasted approximately 4 hours.

### 2.3 Behavioral analysis

Only trials in which participants responded to the primary task, as well as to the interruption task were included in the analysis (incomplete trials older adults: *M* = 33.44, *SD* = 24.73, range = 3 - 102, younger adults: *M* = 9.2, *SD* = 10.56, range = 0 - 42). Trials in which the response was given within the first 150 ms of the response interval (for both the primary task and the interruption task) were excluded from the analysis. Concerning the interruption task, response times and the rate of correct responses were compared between the high-demanding and the low-demanding conditions. For both parameters, a 2 x 2 mixed repeated measures analysis of variance (mixed rm-ANOVA) with *age* (older vs. younger adults) as between-subjects factor and *condition* (high-demanding vs. low-demanding interruption) as within-subject factor was performed.

For the primary task, the difference between the adjusted orientation and the target orientation (angular error) and the time to response initiation (interval between probe presentation and mouse movement onset) was analyzed. The behavioral performance in the primary task was analyzed with a 2 x 3 mixed rm-ANOVA with *age* (older vs. younger adults) as between-subjects factor and *condition* (prolonged fixation vs. high-demanding vs. low-demanding interruption) as a within-subject factor. Additionally, the angular error was compared between the prolonged fixation condition and the early probe condition with a rm-ANOVA with age as between-subjects factor.

### 2.4 EEG recordings, preprocessing and, analysis

Electrophysiological data were recorded with 64 Ag/AgCl passive electrodes (Easycap, GmbH, Herrsching, Germany) in an extended 10/20 scalp configuration (Pivik et al. 1993). Data were sampled by a NeurOne Tesal AC-amplifier (Bittium, Biosignals Ltd, Kuopio, Finland) with a 250 Hz low-pass filter. Impedances were kept below 10 kΩ. The FCz electrode served as reference and the AFz as ground electrode during recording.

Data were analyzed with MATLAB (R2019b) and the EEGLAB toolbox (Delorme & Makeig, 2004) and filtered with a bandpass butterworth filter (pop_basicfilter; low-pass filter with a cutoff frequency of 0.1 Hz, high-pass filter with a cutoff frequency of 30 Hz, filter length = 8 data points). Channels exceeding a kurtosis of 10 SD were rejected (older adults: *M* = 3,32, *SD* = 2,14, range: 0 - 9; younger adults: *M* = 3.88, *SD* = 1.82, range: 0 - 7) by a channel rejection procedure implemented in EEGLAB. After re-referencing the data to common average, the data were segmented into epochs from 700 ms prior to 7500 ms after memory array onset. Trials containing artifacts (threshold = 500 μV, probability threshold = 5 SD, max. % of total rejected trials per iteration = 5%) were excluded based on an automated trial rejection procedure.

Data were down-sampled to 250 Hz to reduce computation time and filtered again with a 1 Hz high-pass filter prior to independent component analysis (ICA). Independent components (IC) associated with eye movements, eye blinks, and generic data discontinuities were identified by ADJUST and rejected. Subsequently, the IC weights were back-projected onto the original low-pass and high-pass filtered 1000 Hz data (0.1 Hz high-pass, 30 Hz low-pass filter). Single dipoles were fitted on the ICs by the EEGLAB plug-in DIPFIT. ICs identified by ADJUST and with a residual variance of the dipole solution exceeding 50% were excluded (remaining ICs older adults: *M* = 78.5%, *SD* = 6.54%; younger adults: *M* = 77.8%, *SD* = 6.57%). The continuous EEG data were again segmented into epochs ranging from 700 ms before to 7500 ms after memory array presentation. The 200 ms interval preceding the memory array was set as the baseline.

The automatic trial rejection procedure, run on the pruned data sets (threshold limit: 1000, probability threshold: 5 SD, max % of trials rejected per iteration: 5%), led-to the exclusion of 106.96 trials on average (range = 53 - 156, *SD* = 31.62) in the sample of older adults and 106.5 trials (range: 61 - 177, *SD* = 36.06) in the sample of younger adults. Only trials with a response to the interruptions task (for the low- and high-demanding interruption conditions) and the primary task were included in the following analyses.

#### 2.4.1 Time-frequency analysis

Event-related-spectral-perturbations (ERSPs) were computed convolving 3-cycle complex Morlet wavelets with each data epoch. Frequencies ranged from 4 to 30 Hz in 52 logarithmic steps and the wavelet cycles increased linearly as a function of frequency with a factor of 0.5, resulting in a 3-cycle wavelet for the lowest frequency and a 11.25-cycle wavelet for the highest frequency. Four hundred time points per epoch were extracted from 282 before to 7082 after memory array onset.

The interval before the memory array (−200 to 0 ms) was taken as spectral baseline for the analysis of the ERSPs. Based on previous research Zickerick et al. (2021), a posterior electrode cluster was used to analyze alpha power (8 - 14 Hz; PO7, PO8, PO3, PO4, POz, P1, P2, P3, P4, P5, P6, P7, P8; see also Erickson et al., 2019)). An electrode cluster surrounding FCz (FC1, FCz, FC2, Fz, Cz; see also Ferreira et al., 2019)) was considered for the analysis of mid-frontal theta power (4 - 7 Hz).

For statistical analysis, the average of both interruption conditions across age groups was compared to the prolonged fixation condition based on a cluster-based permutation procedure. Both posterior alpha and mid-frontal theta power could only be unambiguously related to the re-focusing on the primary task following retro-cue presentation. Thus, we only considered data from the time window between retro-cue onset and memory probe onset for this analysis. Condition labels (interruption vs. no-interruption) were randomly assigned per dataset within 1000 permutations. For each data point, a two-sided within-subject *t*-test (interruption vs. no-interruption) was run for each permutation. This resulted in a time points (37) x permutations (1000) matrix of *p*-values. The size of the largest cluster with *p*-values <.05 was assessed for each permutation. Differences between interruption and no-interruption trials were considered significant if the size of a *p*-value cluster <.05 in the original data was larger than the 95^th^ percentile of the permutation-based distribution of maximum cluster sizes. Subsequently, 2 x 3 mixed analyses of variance (ANOVA) with *age* (older vs. younger adults) as between-subjects factor and *condition* (high-demanding vs. low-demanding interruption vs. prolonged fixation condition) as within-subject factor were performed on the identified time windows to test for an interaction between *age* and *condition*. Since the cluster-based permutation procedure already revealed differences between trials with and without interruptions on EEG level, only *condition x age* interaction effects were considered in the EEG analysis. When a significant interaction was found, a post-hoc ANOVA with *condition* as within-subject factor was performed separately for each age group. In case of a condition main effect, the conditions were additionally compared to each other by follow-up t-tests.

Please note that also the contralateral delay activity (CDA) was computed from a posterior cluster of electrodes contra- and ipsilateral to the relevant information in the memory array (PO7/8, PO3/4, P7/8, and P5/6). This was done to capture the influence of age on the storage of working memory representations during and after the interruption task (Feldmann-Wüstefeld et al., 2018; Vogel & Machizawa, 2004). The respective methods and results are provided as a supplementary section to this manuscript.

#### 2.4.2 Connectivity analysis

The connectivity analysis was performed on trial wise data. Therefore, the time-frequency decomposition was repeated with a custom written code for extracting only the theta frequency range separately for each trial (see Arnau et al., 2019). EEG data of each of the 8200 ms long trials from all channels were convolved with a complex Morlet wavelet, computed as a sine wave in the theta frequency range (center frequency = 4.5) tapered by a Gaussian. The tapering Gaussian was designed with a full width at half maximum (FWHM, Cohen, 2019) of 2.75 in the frequency domain and 298 in the time domain.

Functional connectivity was analyzed by computing the phase-lag index (PLI, Stam et al., 2007). The PLI assesses an asymmetry of non-zero and non-π valued phase differences across trials at a given point in time. When 50% of all non-zero phase values between two time-series (here we only compared two theta ranges) of all trials range between – π and 0 and the other 50% range between 0 and π, the PLI has a value of zero. Functional connectivity between two sides of electrodes is indicated by an asymmetry in the distribution of phase values. Thus, the PLI equals one, when all difference values are in one of these two ranges. Since the PLI is insensitive to the distribution of phase value differences of 0 or π, it is robust to sources represented in both time series due to volume conduction.

The PLI was computed to complement the analysis of oscillatory power in the theta frequency range. Thus, the connectivity was computed between each electrode of the mid-frontal cluster surrounding FCz (FC1, Fz, Cz, FCz, FC2) and all other electrodes. Afterwards, the PLI values were averaged across the five electrodes for building a cluster of seed electrodes. The connectivity analysis was performed on the time window revealed by the cluster-based permutation procedure for the theta frequency range as described above (91-516 ms after retro-cue onset). Within this time window, we calculated the PLI values averaged across experimental conditions separately for each age group. Only electrodes with a PLI to the seed cluster exceeding the average PLI across all electrodes by at least one standard deviation were included in the further analysis (older adults: O2, Pz, PO10, O10, P2, PO5, P6 CPz, and POz; younger adults: O1, O2, Oz, PO10, O9, O10, PO4, POz). The PLI values were compared by a mixed rm-ANOVA with *condition* (high-demanding vs. low-demanding interruption vs. prolonged fixation condition) as within-subject and *age* (older vs. younger adults) as between-subjects factor.

### 2.5 Inferential statistics and effect sizes

Results of the neuropsychological tests were compared between age groups with *t-*tests for independent samples. In cases were the variance differed between the age groups, Welch’s test was used to compare the performance in the neuropsychological tests between the age groups.

For all ANOVAs, Greenhouse-Geißer correction was applied when sphericity, as assessed with the Mauchly tests for sphericity, was violated (indicated by *ε*). To prevent the effects of interest from *p*-value inflation due to multiple testing within each ANOVA, FDR correction was applied whenever more than one factor was analyzed. We reported adjusted p-values (*p_adj_*) in this regard (Cramer et al., 2016). Furthermore, FDR correction was also applied for multiple testing in all post-hoc comparisons (*p_adj_*; see Benjamini & Hochberg, 1995).

## 3 Results

### 3.1 Behavioral data

#### 3.1.1 Interruption Task

A significant interaction between *age* and *condition* was found on the level of response times, *F(1,41)* = 10.52, *p* = .003, *η^2^_p_* = .20. In both age groups, participants required more time to respond to a high-demanding interruption task. However, the difference in response time between the interruption conditions was weaker in older than in younger adults *(M)_(older)_* = 1435 ms, *SD_(older)_* = 99.06; *M)_(younger)_* 1263 ms, *SD_(younger)_* 120.17; *M_(older)_* = 1180 ms, *SD_(older)_* = 148.28; *M_(yonger)_* = 877.16, *SD_(younger)_* = 146.49; older adults: *t(17)* = 10.29, *p_adj_* < .001, *d_z_* = 2.43; younger adults: *t(24)* = 14.02,*p_adj_* < .001, *d_z_* = 2.80, 95% CI [220.36, 312.59]).

Percentage of correct responses was lower in the high-demanding interruption task than in the low-demanding interruption task, *F(1,41)* = 42.36, *p_adj_* < .001, *η^2^_p_* = .51 (*M_(high)_* = 8.00, *SD_(high)_* = .14; *M_(low)_* = 0.92, *SD_(low)_* = .10). Additionally, the rate of correct responses was lower for the older (*M*= 0.82, *SD* = 0.13) than for the younger age group (*M*= 0.90, *SD* = 0.07), *F(1,41)* = 7.16,*p_adj_* = .016, *η^2^_p_* =.15. There was no *age x condition* interaction, *F*(41,1) = 0.04, *p_adj_* = .845, *η^2^_p_* <.001.

#### 3.1.2 Primary Task

For assessing performance in the primary task, the time to initialize the response (time to mouse movement onset, see figure 2A), as well as the difference between the adjusted probe bar and the target orientation in degree (angular error, see figure 2B), were analyzed. All analyses were based solely on trials in which subjects had confirmed the completion of orientation adjustment by button press and had completed the interruption task.

**Figure 2.**
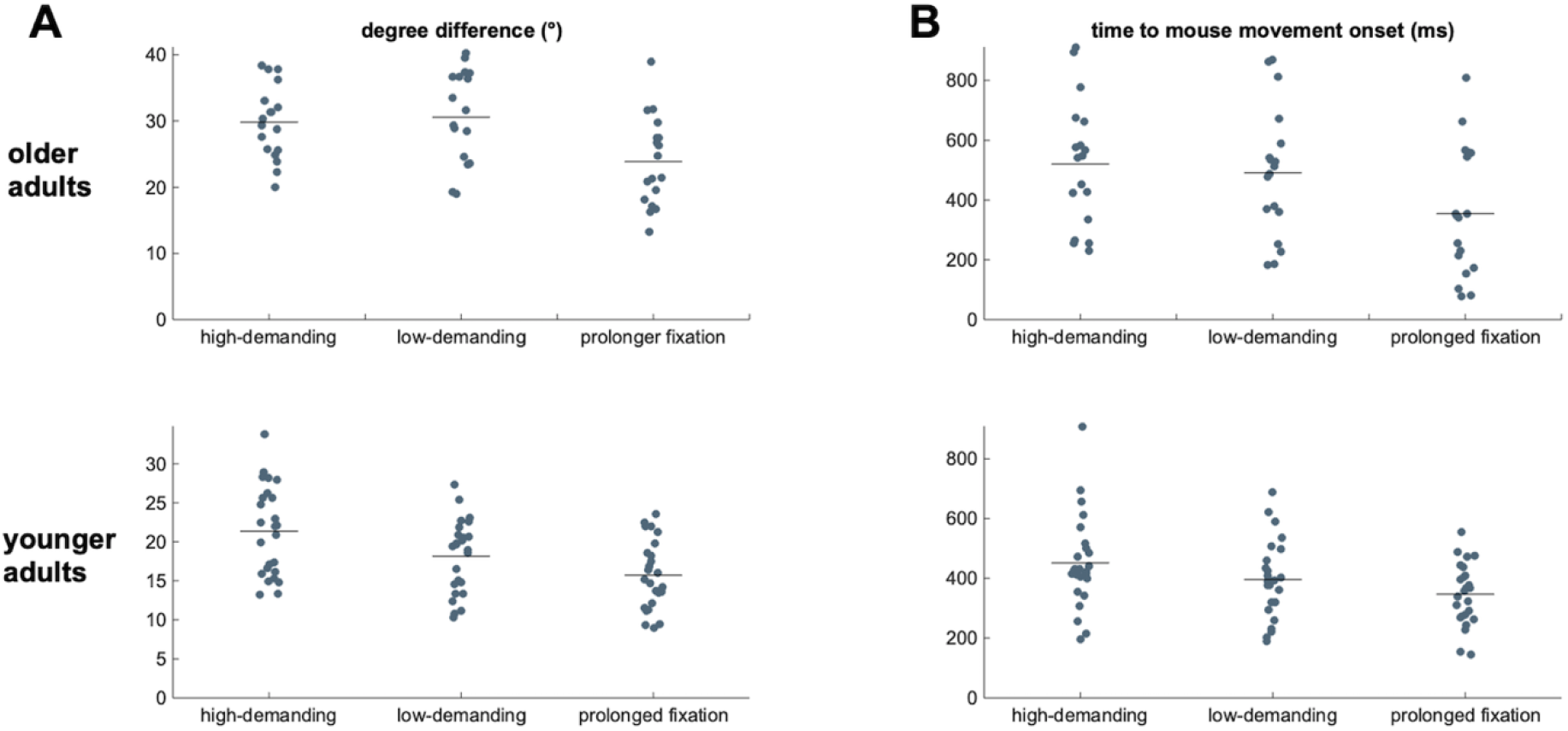
Behavioral results. 2A depicts the raw angular error for each age group separately for the high-demanding and low-demanding interruption conditions and the prolonged fixation condition. The vertical line indicates the condition average within each age group, while each dot represents the mean angular error of one participant. The time to mouse movement onset (in ms) for each condition and age group is shown in 2B.

Analysis of time to mouse movement onset revealed a difference between age groups, *F(1,41)* = 5.29, *p_adj_* = .048, *η^2^_p_* = 0.11, ε = .63, and between conditions, *F(2,82)* = 266.39, *p_adj_* < .001, *η^2^_p_* = 0.86, ε = .63 (see figure 2). The interaction effect reached significance as well, *F(2,82)* = 3.75, *p_adj_* = .048, *η^2^_p_* = .08, ε = .63. Post-hoc ANOVAs revealed a modulation of time to mouse movement onset by condition in both age groups (older adults: *F(2,34)* = 22.15,*p_adj_* < .001, *η^2^_p_* = 0.57; younger adults: *F(2,48)* = 27.38,*p_adj_* < .001, *η^2^_p_*= 0.53). Interestingly, there was no difference in time to mouse movement onset between the interruption conditions for older adults, *t(17)*= 1.98, *p_adj_* = .246, *d_z_* = 0.47, CI 95% [-2.87, 89.1] (*M_(high)_* = 563.63, *SD_(high)_* = 215.96; *M_(low)_* = 520.52, *SD_(low)_* = 216.34), while younger adults needed significantly more time to initialize the response after a high-demanding (*M_(high)_* = 450.94, *SD_(high)_* = 154.35) than after a low-demanding interruption (*M_(low)_* = 395.96, *SD_(low)_* = 129.65), *t(24)* = 4.47, *p* < .001, *d_z_* = 0.89, CI 95% [29.62, 80.34].

The angular error revealed a significant *condition x age* interaction, *F(2,82)* = 8.13, *p* < .001, *η^2^_p_* =.17. The condition effect was weaker in the older than the younger age group, as demonstrated by separate post-hoc ANOVAs for each age group (older adults: *F(2,34)* = 23.81, *p*_adj_ < .001, *η^2^_p_* = .58; younger adults: *F(2,48)* = 46.04, *p_adi_* < .001, *η^2^_p_* = .66, ε = .77). Both interruption conditions increased the angular error in the older age group to a similar extent (*M_(high)_* = 30.60, *SD_(high)_* = 5.74; *M_(low)_* = 31.12, *SD_(low)_* = 6.35; high-demanding interruption vs. prolonged fixation: *t(17)* = −5.62, *p_adj_* < .001, *d_z_* = −1.32, CI 95%[−8.38, −3.81], low-demanding interruption vs. prolonged fixation: *t(17)* = −5.63, *p_adj_* < .001, *d_z_* = −1.33, CI 95%[−9.09, −4.13], high-demanding vs. low-demanding interruption: *t(17)* = −0.56, *p* = .581, d*z* = −0.13, 95% CI [-2.47, 1.43]). For younger adults, the angular error was increased by a low-demanding interruption compared to the prolonged fixation condition (*M_(fix)_* = 15.71, *SD_(fix)_* = 4.40; *M_(low)_* = 18.15, *SD_(low)_* = 4.68), *t(24)* = −5,75, *p_adj_* < .001, *d_z_* = −1.15, CI 95% [−3.31, − 1.56]) and even more increased following a high-demanding interruption (*M_(high)_* = 21.37, *SD_(high)_* = 5.74; *t(24)* = 5.40, *p_adj_* < .001, *d_z_* = 1.15, CI 95% [1.99, 4.45]).

It might be that the time available for rehearsing the working memory items increased primary task performance in the prolonged fixation condition, for example by the application of strategies for transforming the visuo-spatial representation into verbal codes following encoding. In this case, the relative decrease in primary task performance in the interruption conditions compared to the prolonged fixation condition would not be unambiguously related to interference by the interruption task. To rule out this possibility, a mixed ANOVA with *age* as between-subjects factor and the *no-interruption conditions* (early probe condition vs. prolonged fixation condition) as within-subject factor revealed a main effect of condition on task performance, *F(2,41)* = 18.87, *p* = .011, *η^2^_p_* = .15. The angular error was even higher in the prolonged fixation condition (*M* = 19.40, *SD* = 6.78) compared to the early probe condition (*M* = 18.42, *SD* = 6.48), *t(42)* = 2.80, *p* = .008, *d_z_* = 0.43, CI 95% [0.27, 1.68]). Therefore, rehearsal strategies possibly used in the prolonged retention interval cannot have led to an increase in performance compared to the interruption conditions.

### 3.2 EEG data

#### 3.2.1 Posterior alpha power

For posterior alpha power, the cluster-based permutation procedure revealed a difference between the interruption conditions and the prolonged fixation condition 294 – 682 ms following the retro-cue (see Fig. 3). There was also a *condition x age* interaction, *F(2,82)* = 4.35, *p*= 0.021, *η^2^_p_* =.1, ε = .85. Subsequent analyses revealed a strong condition effect for the older adults, *F(2,34)* = 15.81, *p_adj_* < .001, *η^2^_p_* = .48, ε = .72, with a lower alpha suppression following interruptions. Relative to this, the younger adults showed a weaker condition effect, *F(2,48)* = 3.93, *p_adj_* = .03, *η^2^_p_* = .14.

**Figure 3.**
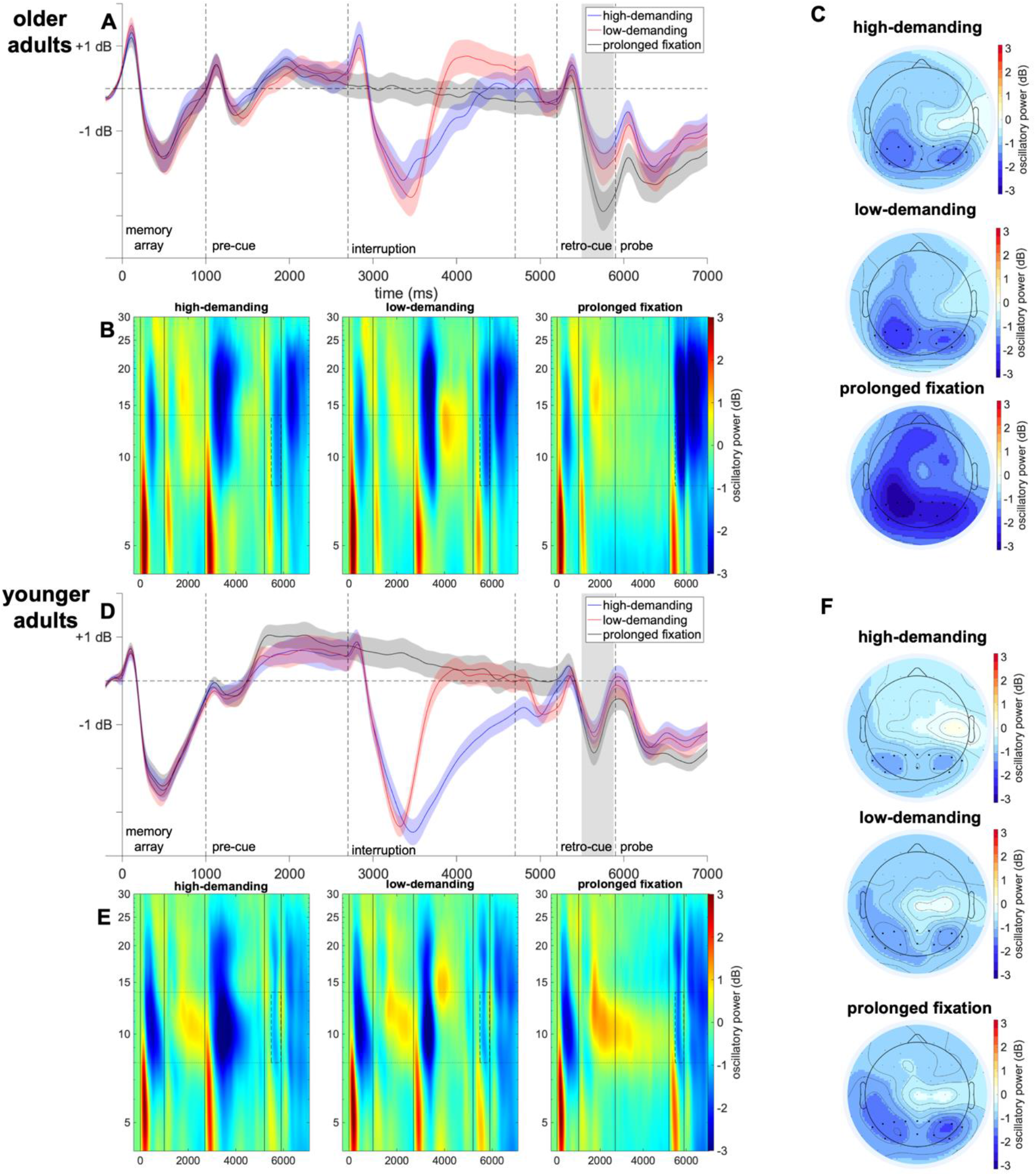
Posterior alpha power. ERSPs from a cluster of posterior channels in the alpha frequency range are depicted over the whole trial separately for older (3A and 3B) and for younger adults (3D and 3E). Events in the trial are indicated by vertical lines (0 ms memory array on-set; 1000 ms pre-cue onset; 2700 ms interruption onset; 5200 ms retro-cue onset; 5900 ms memory probe onset). The time window from 294 to 682 ms following the retro-cue was revealed by a cluster-based permutation procedure and is highlighted with grey areas. Figures 3C (older adults) and 3F (younger adults) depict the topographical distribution of alpha power for the interruption conditions and the prolonged fixation condition.

Post-hoc *t*-tests revealed that alpha power suppression following retro-cue presentation was reduced by a preceding interruption in both age groups. The alpha power suppression was significantly weaker after the high-demanding interruption than the prolonged fixation period (older adults: *t(17)* = 4.36, *p_adj_* = .003, *d_z_* = 1.03, 95% CI [.06, 1.66]; younger adults: *t(24)* = 2.28, *p_adj_* = .047, *d_z_* = 0.46, 95% CI [0.04, 0.76]). A similar pattern was revealed for the low-demanding interruption and the prolonged fixation condition (older adults: *t(17)* = 4.04, *p_adj_* = .003, *d_z_* = 0.95, 95% CI [0.42, 1.33]; younger adults: *t(24)* = 2.65, *p_adj_* = .028, *d_z_* = 0.53, 95% CI [0.42, 1.33]). However, in neither age group was alpha power suppression affected by the type of interruption (older adults: *t(17)* = 1.79, *p_adj_* = .109, *d_z_* = 0.42, 95% CI [-0.04, 0.53]; younger adults: *t(24)* = 0.45, *p_adj_* = .656, *d_z_* = 0.09, 95% CI [-0.25, 0.39]).

#### 3.2.2. Mid-frontal theta power

Mid-frontal theta oscillations are depicted in Figure 4. The cluster-based permutation procedure revealed significantly lower theta power following the interruption conditions than in the prolonged fixation condition 91-516 ms after retro-cue presentation. No significant *age x condition* interaction was found in this time window, *F(2,82)* = 0.59, *p*= 0.522, *η^2^_p_* =.01, ε = .81. Thus, post-hoc analyses were performed on the whole sample. Following a high-demanding interruption, theta power was even lower than following a low-demanding interruption, *t(42)* = −4.04, *p_adj_* < .001, *d_z_* = −0.62, 95% CI [-0.59, −0.20].

**Figure 4.**
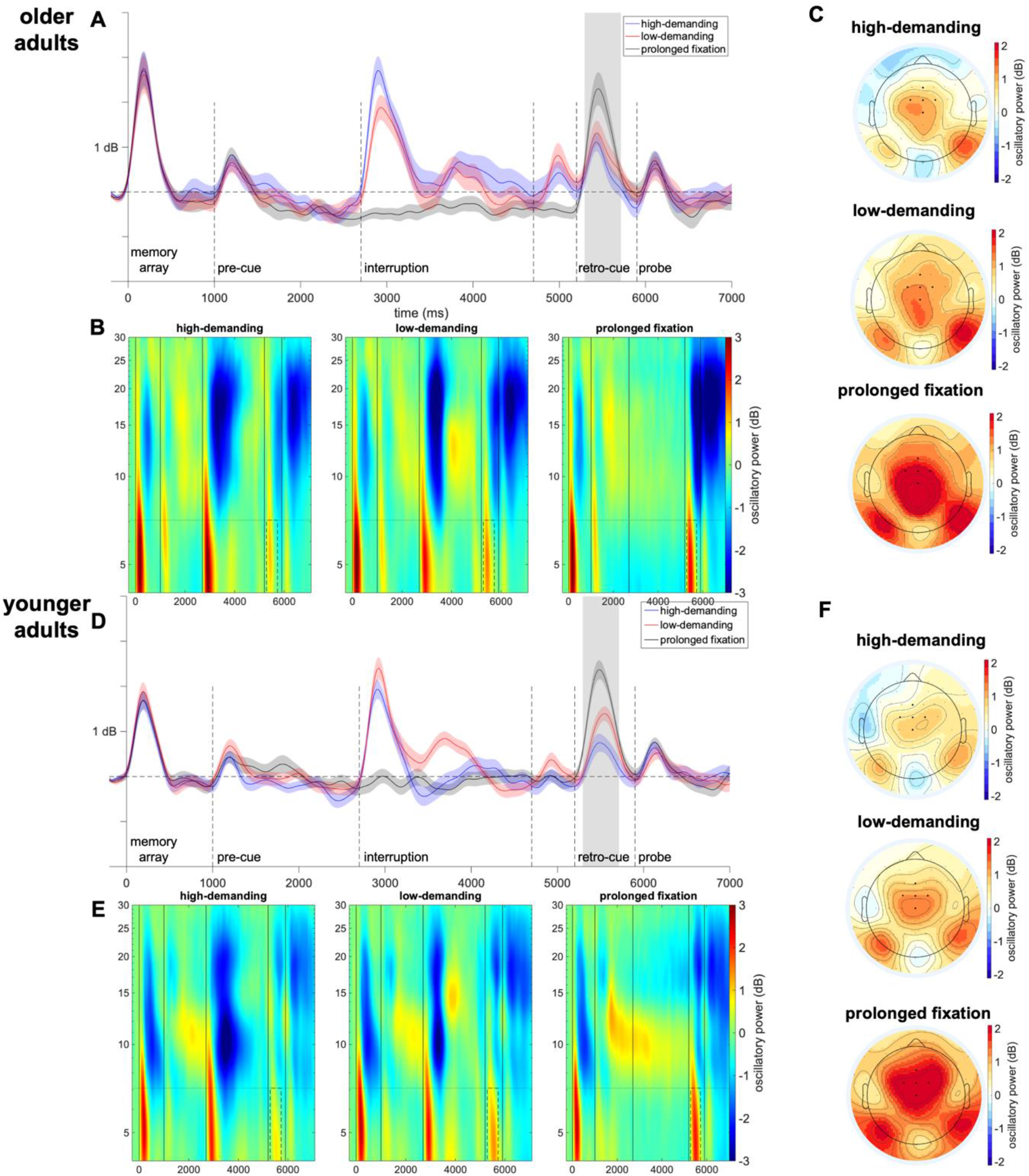
Frontal theta power. Oscillatory power at mid-frontal electrodes in the theta-frequency range is depicted separately for each condition and each age group (older adults: 4A and B, younger adults: 4D and E). The vertical lines indicate the events within a trial (0 ms memory array onset, 1000 ms pre-cue onset, 2700 ms interruption onset, 4700 interruption offset, 5200 ms retro-cue onset, 5900 ms memory probe onset). A time window 91-516 ms after the retro-cue was revealed by a cluster-based permutation procedure and is highlighted with grey areas. The differences between each interruption and the prolonged fixation condition are depicted on the right side. Figures 4C (older adults) and 4F (younger adults) depict the topographical distribution of alpha power for the interruption conditions and the prolonged fixation condition.

#### 3.2.3 Connectivity analysis

To complement the theta power analysis, connectivity in the theta frequency range was analyzed between mid-frontal and electrodes exceeding the average connectivity values by at least one standard deviation (see figure 5; electrode cluster older adults: O2, Pz, Oz, PO10, O10, P2, PO4, P6, CPz, POz; electrode cluster younger adults: O1, O2, Oz, PO10, O9, O10, PO4, PO). A main effect of *age, F(1,41)* = 4.19, *p* = 0.047, *η^2^_p_* =.09, ε = .86, and a respective interaction was found in the connectivity analysis, *F(2,82)* = 4.60, *p* = 0.013, *η^2^_p_* =.10, ε = .86. Post-hoc analysis revealed that there was no modulation by condition of PLI values for older adults, *F(2,34)* = 1.76, *p_adj_* = .187, *η^2^_p_* = .09. However, for younger adults, phase coherence between mid-frontal and posterior electrodes differed between the conditions, *F(2,34)* = 13.75, *p_adj_* < .001 *η^2^_p_* = .36. It was weakened by a preceding high-demanding interruption relative to the prolonged fixation condition, *t(24)=* −3.66, *p_adj_* = .001, *d_z_* = −0.73, 95% CI [−0.06, - 0.02], but not by a preceding low-demanding interruption, *t(24)=* .49, *p_adj_* = .625, *d_z_* = 0.05, 95% CI [-0.01, 0.02]. Theta connectivity was also lower following a high demanding interruption than following a low demanding one, *t(24)=* −5.74, *p_adj_* < .001, *d_z_* = 1.15, 95% CI [0.03, 0.06].

**Figure 5.**
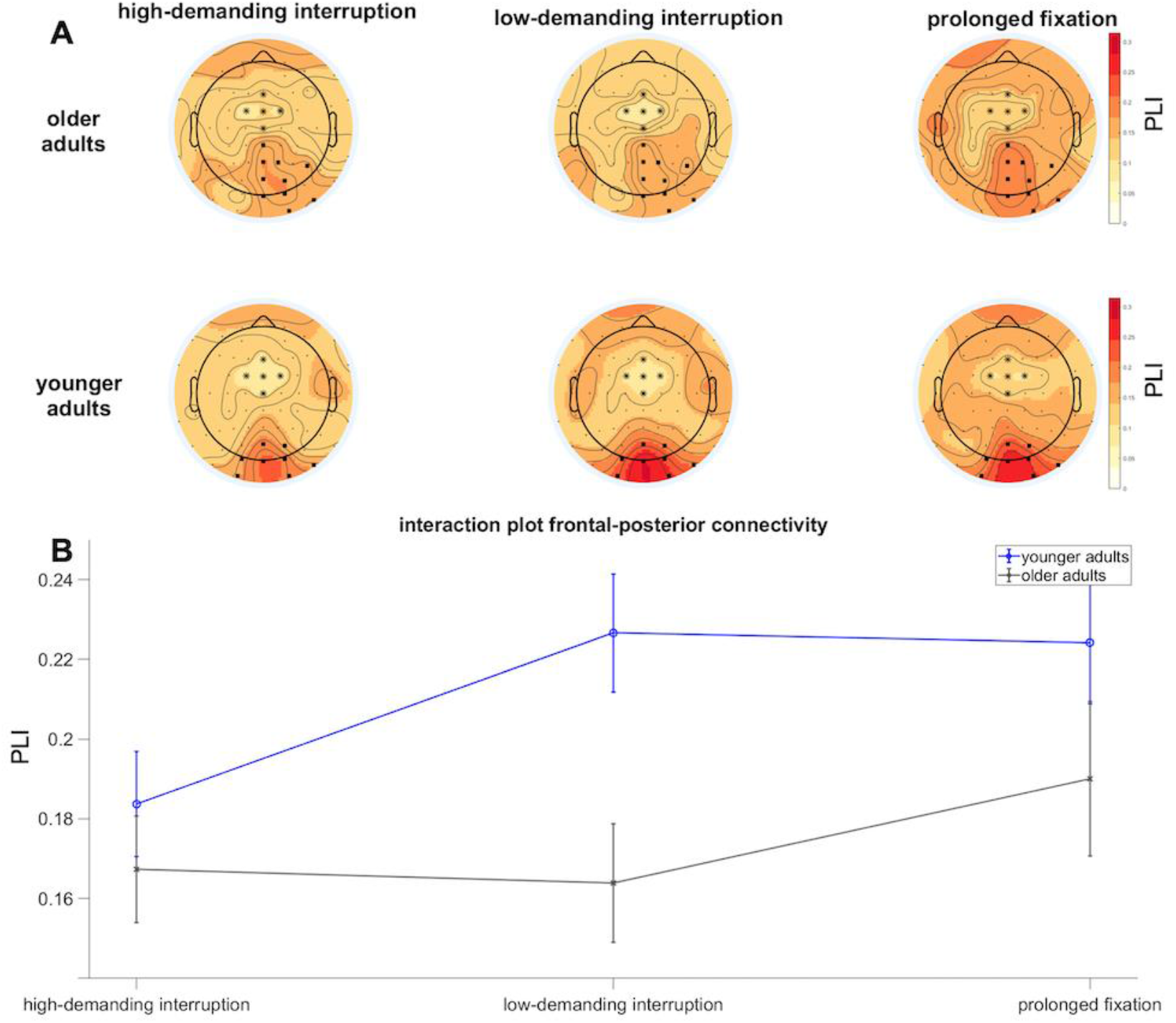
Phase lag index. The PLI (5A) 91-516 ms after retro-cue onset was computed between a seed cluster of mid-frontal electrodes (FCz, Fz, Cz, FC1, FC2; marked with stars) and the O2, Pz, Oz, PO10, O10, P2, PO4, P6, CPz and POz electrodes for older and the O1, O2, Oz, PO10, O9, O10, PO4 and POz electrodes for younger adults (marked with squares). Panel B depicts the interaction of age group and condition on frontal-posterior connectivity.

## 4 Discussion

The present study investigated the effects of aging on primary task resumption after an interruption. To this end, a working memory experiment was conducted requiring participants to memorize the orientation of two bars. At the end of each trial, participants had to reproduce the orientation indicated by a retro-cue. Importantly, prior to retro-cue presentation, the task was interrupted by either a high- or low-demanding task. This way we could demonstrate that aging decreases the ability to select primary task information after an interruption.

On behavioral level, we could show that the detrimental effect of interruptions on task performance (accuracy and time to movement onset) was pronounced when the interruption was cognitively high-demanding compared to when it was low-demanding (see also Cades et al., 2008; Gillie & Broadbent, 1989; Hodgetts & Jones, 2006; Zickerick, Rösner, et al., 2021), but only for younger adults. Primary task performance in older adults was independent of the cognitive requirements of the interruption task. Still, for both age groups, behavioral data from the interruption task showed that the high-demanding interruption task was indeed more difficult to perform than the low-demanding interruption task. It is nevertheless possible that the successful completion of the low-demanding interruption task also required the complete withdrawal of attentional resources from the primary task in older adults, which could cause the performance deficit to no longer differ between the two interruption conditions. This is also in line with the observation that the performance difference between the high- and low-demanding interruption task was weaker for the older adults. The age-related effect on primary task performance might further be explained by a floor effect for older adults. The presence of an interruption might already have decreased working memory performance in older adults to such an extent, that even cognitively more demanding interruptions did not induce an additional performance deficit. However, these two explanations cannot be specifically tested on the basis of the current experiment.

The main goal of this study was to understand the effect of aging on neurocognitive processes underlying primary task resumption. The retro-cue required participants to refocus attention only on the target representation or, in other words, to selectively attend to only one primary task item. Thus, oscillations in the theta and alpha frequency range following the retro-cue were analyzed as correlates of (retrospective) attentional control processes. Furthermore, as a marker for active storage of visuo-spatial information in working memory, the contralateral delay activity (CDA; Vogel & Machizawa, 2004; see supplementary information) was analyzed. CDA amplitude at the end of the interruption interval was not affected by the presence or absence of an interruption nor did it differ between age groups. Thus, even though interruptions required participants to shift the focus of attention between the primary and the interruption task, working memory storage remained stable across conditions and age groups.

Still, in line with our previous study, a preceding interruption impaired the ability to flexibly apply attentional control resources on the level of working memory (Zickerick et al., 2021). This was the case in both age groups, reflected in lower mid-frontal theta power after the retro-cue when it was preceded by a task interruption. Even though older adults showed no difference on a behavioral level between working memory performance after a high- and a low-demanding interruption, available control resources in both age groups were diminished by a high-demanding compared to a low-demanding interruption. Contrary to previous findings suggesting that aging decreases executive functioning (Cummins & Finnigan, 2007; Gazzaley et al., 2007; Wascher et al., 2012), we found no difference in the effect of interruptions on theta power following the retro-cue between the age groups. The dual mechanisms of control framework (Braver, 2012; Braver et al., 2008) can reconcile this discrepancy by dividing the concept of cognitive control into proactive and reactive processes. Proactive control is a form of early selection in which task-relevant information is actively maintained, while reactive control rather functions as a late correction mechanism, e.g., is active after interference detection or, in the case of this experiment, following the retro-cue. Thus, proactive control is resource consuming compared to reactive control. In line with this, it has been shown that older adults rely more on reactive control processes than younger adults (Arnau et al., 2019; Braver, 2012; Braver et al., 2001). This stronger use of reactive control processes might explain the lack of age-related differences regarding the effect of interruptions on theta power levels following the retro-cues.

However, although mid-frontal theta power following the retro-cues did not reveal any age-effects, long-range functional connectivity in the theta band did. Mid-frontal seed electrodes exhibited the strongest connectivity to posterior electrodes. The posterior clusters differed between the older and younger age groups: the electrodes were slightly more posterior in younger compared to older adults. However, since this analysis was performed on sensor level and not on source level, we cannot draw strong conclusions from this difference (Palva & Palva, 2012). Nevertheless, the effect of interruptions on functional connectivity regardless of the electrode clusters differed between the age groups. Theta connectivity was modulated by the demands of the interruption task only for younger adults. It was higher in the low-demanding interruption and prolonged fixation conditions than following high-demanding interruptions. Tóth et al. (2014) related frontal-midline theta connectivity to parietal and occipital areas to the refreshing of information during working memory storage. They could show that connectivity was reduced in older adults and correlated to memory performance irrespective of age. The current results also show generally lower theta connectivity in older adults (see figure 5). Thus, while younger adults are able to selectively maintain the relevant primary task information in working memory, at least after a low-demanding interruption, the connectivity patterns for older adults indicate a general deficiency in this regard. This might also explain the findings at the behavioral level that older adults’ working memory performance was equally affected by interruption tasks of varying difficulty.

Furthermore, posterior alpha power suppression after the retro-cue was decreased by a preceding interruption in both age groups. This replicates our previous finding that interruptions in general impair primary task resumption (Zickerick et al., 2021). Suppression of oscillatory power in the alpha frequency range over posterior electrodes has been linked to attentional selection of cued working memory content (Hajonides et al., 2019; Schneider et al., 2017). Older adults showed a generally stronger suppression of alpha power after the retrocue, which might indicate the need for more attentional resources to select the relevant primary task information. Importantly, the effect of interruptions on this alpha suppression was stronger for the older participants, indicating a deficit in the focusing of attention on primary task information following an interruption. This could, for example, be due to the fact that older adults found it more difficult to withdraw attention from the then irrelevant interruption task, which would be in line with previous research pointing to an age-related deficit in the handling of irrelevant information (see Gazzaley et al., 2008; Mertes et al., 2017).

In summary, we showed that both older and younger adults were impaired in their working memory performance by an interruption. However, only younger adults showed a deficit as a function of the difficulty of the interruption task. On the one hand, this can be explained by a general age-related deficit in actively maintaining relevant information in working memory, indicated by reduced theta connectivity between fronto-central and posterior brain regions. On the other hand, older adults also showed a greater attenuation in the suppression of posterior alpha power following the task interruptions, which shows that they were less able to cope with the negative influences of getting distracted (by the interruptions) on the selection of relevant primary task information.

## Acknowledgements

The authors would like to thank Tobias Blanke for programming the experiment, Pia Deltenre, Barbara Foschi and Katrine Bergeron for assistance in collecting the data, and Andre Lauff und Asli Yavuz for evaluation of the neurophysiological tests, and Kira Dolhan for prove reading the manuscript.

## Supplementary Material

### Contralateral delay activity

For the analysis of the contralateral delay activity (CDA) the event-related potential was averaged across two clusters of posterior electrodes ipsilateral and contralateral to the relevant stimuli in the memory array (PO7/8, PO3/4, P7/8, and P5/6). To analyze whether impairment of active storage of primary task visual spatial representations differed between the age groups, contra-minus ipsilateral difference waves were computed separately for age group and condition. The data were further averaged over the 500 ms time window between interruption offset and retro-cue onset (4700 – 5200 ms after memory array onset).

The grand average of all conditions and all participants revealed a strong CDA after the memory array. A within-subjects *t*-test against zero based on the grand average demonstrated that the CDA reappears after interruption offset (4700 ms after the memory array), *t*(42) = −4.64, *p* < .001, d_z_ = −0.71. Thus, after completing interruption processing participants refocus their attention on primary task representations. Importantly, a mixed ANOVA with the within-subjects factor *condition* (high-demanding interruption, low-demanding interruption, prolonged fixation) and the between subjects-factor *age* (older adults, younger adults) revealed neither a difference between conditions (*F*(2,82) = 2.09, *p* = .132, *η^2^_p_* = .05) nor between age groups ( *F*(1,41) = 1.66, *p* = .204, *η^2^_p_* = .04). Furthermore, there was no interaction between the factors, *F*(2,82) = 0.48, *p* = .618, *η^2^_p_* = .01 This demonstrates that the storage of a visual-spatial representation within working memory is neither affected by a preceding interruption nor does it differ between the age groups.

**Supplementary figure S1.**
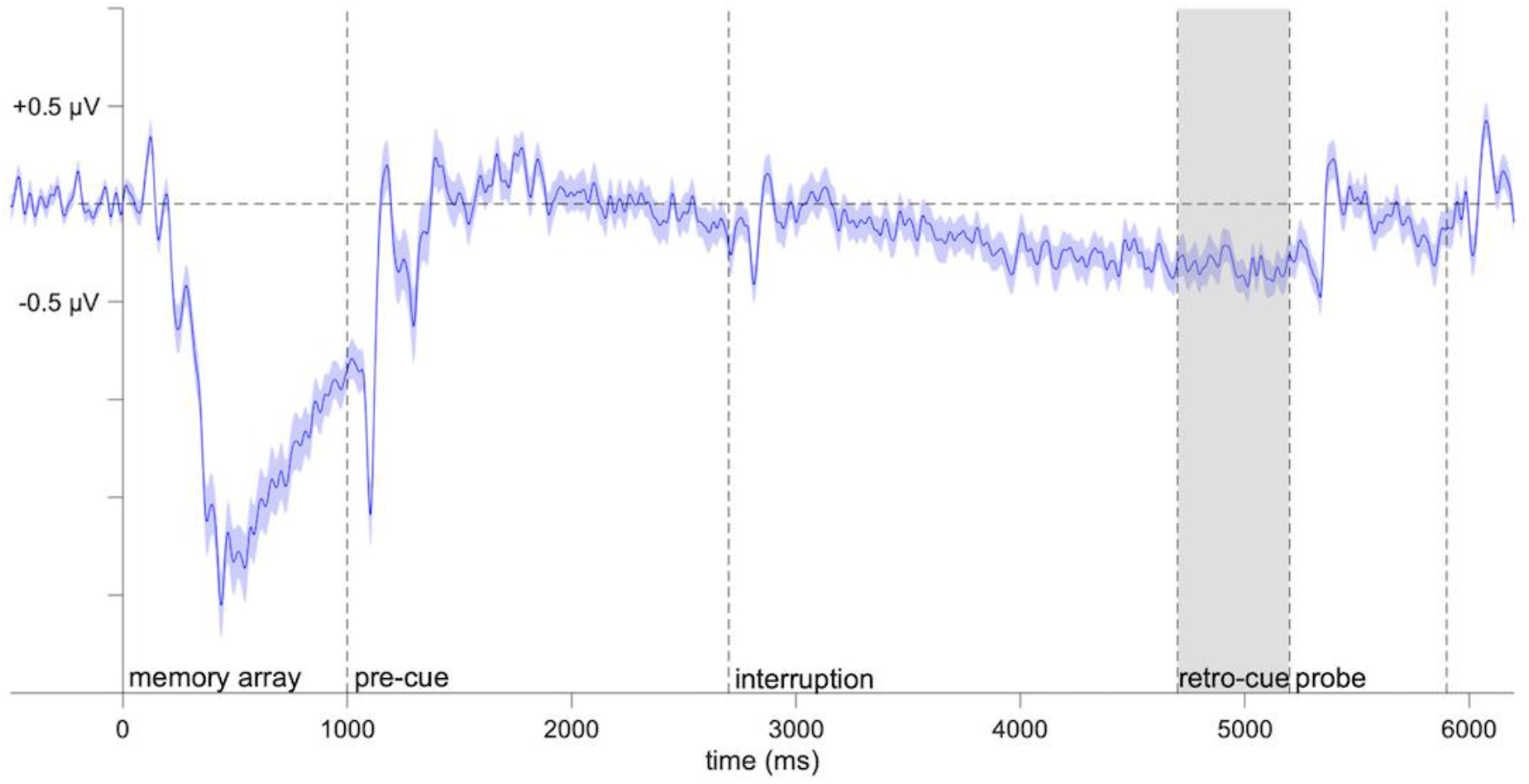
Contralateral delay activity (CDA). Illustrated is the CDA as a difference wave (contra-ipsilateral to target position in the memory array) averaged across all conditions and subjects. The grey areas highlight the analyzed time window (4700 – 5200 ms after memory array onset).

